# Mitochondrial and bioenergetic processes are enriched in large- but are reciprocally depleted in small extracellular vesicles

**DOI:** 10.64898/2026.10.23.761372

**Authors:** Nikhila Achanta, Devika S Manickam

**Author notes:** Corresponding author: Devika S Manickam 6431 Fannin Street, MSE R274 Houston, TX 77030.

## Abstract

Extracellular vesicles (**EVs**) are a promising class of drug delivery carriers for a broad range of therapeutic applications. We have previously demonstrated that brain endothelial cell (**BEC**)-derived large EVs (>200 nm in particle diameter) increased mitochondrial function in recipient BECs and improved therapeutic outcomes in a mouse model of ischemic stroke. In the current study, we demonstrate using size-resolved quantitative proteomics and gene-set enrichment analysis that *mitochondrial and bioenergetic processes are enriched in large- but are reciprocally depleted in small EVs* (exosomes, <200 nm in particle diameter). <u>These results are striking and reveal the selective packaging of mitochondrial cargo in large but not small EVs.</u> These strong mechanistic data anchor the rationale for the continued development of large EV mitochondria as a potential approach for the treatment of multiple diseases associated with mitochondrial dysfunction.

## Introduction

Intercellular transfer of mitochondria is an emerging mechanism of cerebroprotection in the central nervous system, and extracellular vesicles (**EVs**) have been identified as vehicles for this type of transfer.^1-3^ We have previously demonstrated that brain endothelial cell (**BEC)-**derived large EVs (**lEVs**) (particle diameters > 200 nm), but not the smaller EVs: **sEVs**/exosomes, transfer functional mitochondria to recipient BECs and increase recipient ATP under ischemic conditions,^4^ and large EVs delivered to the BBB reduce injury in a mouse model of ischemic stroke.^5, 6^ Whether the mitochondrial cargo of BEC EVs is systematically partitioned by vesicle size at the level of the whole proteome has not been defined. Size-resolved quantitative proteomics were performed on lEVs and sEVs from mouse b.End3 brain endothelial cells and we used gene-set enrichment analysis to map the biological processes enriched in each subpopulation.

## Methods

### EV isolation

Prior to EV isolation, donor mouse brain endothelial cells (b.End3) were maintained in DMEM medium for 48h containing heat-inactivated (**HI**) or non-heat-inactivated (**non-HI**) exosome-depleted fetal bovine serum, giving a 2 × 2 design (EV size × serum). EVs were harvested from conditioned medium and separated into lEV and sEV subpopulations by sequential centrifugation as previously described.^4-6^ Three independent preparations at serial passages (P27-P29) served as biological replicates.

### Proteomics

EV protein samples were denatured in 5% SDS, reduced with 5 mM DTT (90°C, 5 min), and alkylated with 10 mM iodoacetamide (15 min, room temperature). Proteins were processed on S-Trap Micro Spin Columns (ProtiFi) per manufacturer protocol and digested overnight at 37°C with a trypsin/LysC mixture. Peptide concentration was determined by colorimetric assay (Pierce Quantitative Colorimetric Peptide Assay, Thermo Scientific) and samples were reconstituted at 100 ng/µL in 0.1% formic acid containing 0.015% DDM. LC-MS/MS analysis was performed on a nanoElute2 UHPLC system coupled online to a timsTOF Ultra 2 mass spectrometer (Bruker Daltonics) using a 25 cm × 75 µm × 1.5 µm PepSep column with a 28-min active gradient. Data were acquired in diaPASEF mode (m/z 400–1000; ion mobility 0.64–1.37 1/K0; 24 DIA windows; cycle time 0.96 s).

Raw data were searched in library-free mode using DIA-NN (v2.2.0) against a mouse proteome FASTA (UniProtKB; 17,228 entries; accessed November 2024). Search parameters included trypsin/P digestion (up to 1 missed cleavage), variable oxidation (M) and N-terminal acetylation, fixed carbamidomethylation (C), and precursor FDR set at 1% with match-between-runs enabled. Protein quantification was performed using the iBAQ approach via the gpGrouper algorithm, followed by median normalization and log2 transformation.^7^ Differential expression was assessed using moderated t-tests implemented in the R package limma with Benjamini-Hochberg FDR correction.

### Analysis

Differential abundance was assessed with limma moderated t-tests and Benjamini-Hochberg correction (FDR< 0.05, >= 2-fold change). Gene set enrichment analysis (**GSEA**) was performed with fgsea against MSigDB C% GO Biological Process collection in two modes: per sample, ranking proteins by z-scored abundance (one normalized enrichment score, **NES**, per gene set per sample), and per comparison, ranking proteins by signed log p-value from the lEV *vs*. sEV comparisons under each serum condition.^7^ C5 GO Biological Processes were of importance because gene otology and pathway enrichment in our pilot proteomic analysis of BEC-derived EVs resolved their cargo at the level of biological processes, most prominently glycolysis-related processes.^4^ Our prior biochemical and functional studies established EV mitochondria as therapeutic cargo, which directed our attention to mitochondrial and bioenergetic processes.^4-6, 8^ Enrichment was considered significant at adjusted p < 0.25 are marked in figures. Leading -edge proteins are reported for enriched sets.

## Results

In total, 4,981 proteins were quantified. Unsupervised hierarchical clustering of per-sample NES profiles partitioned the twelve samples primarily first by EV size, with serum condition and passage sorting only within the lEV and sEV branches (**Figure 1A**). Vesicle size, rather than serum treatment or passage number, was therefore the main source of variation in EV cargo.

**Figure 1.**
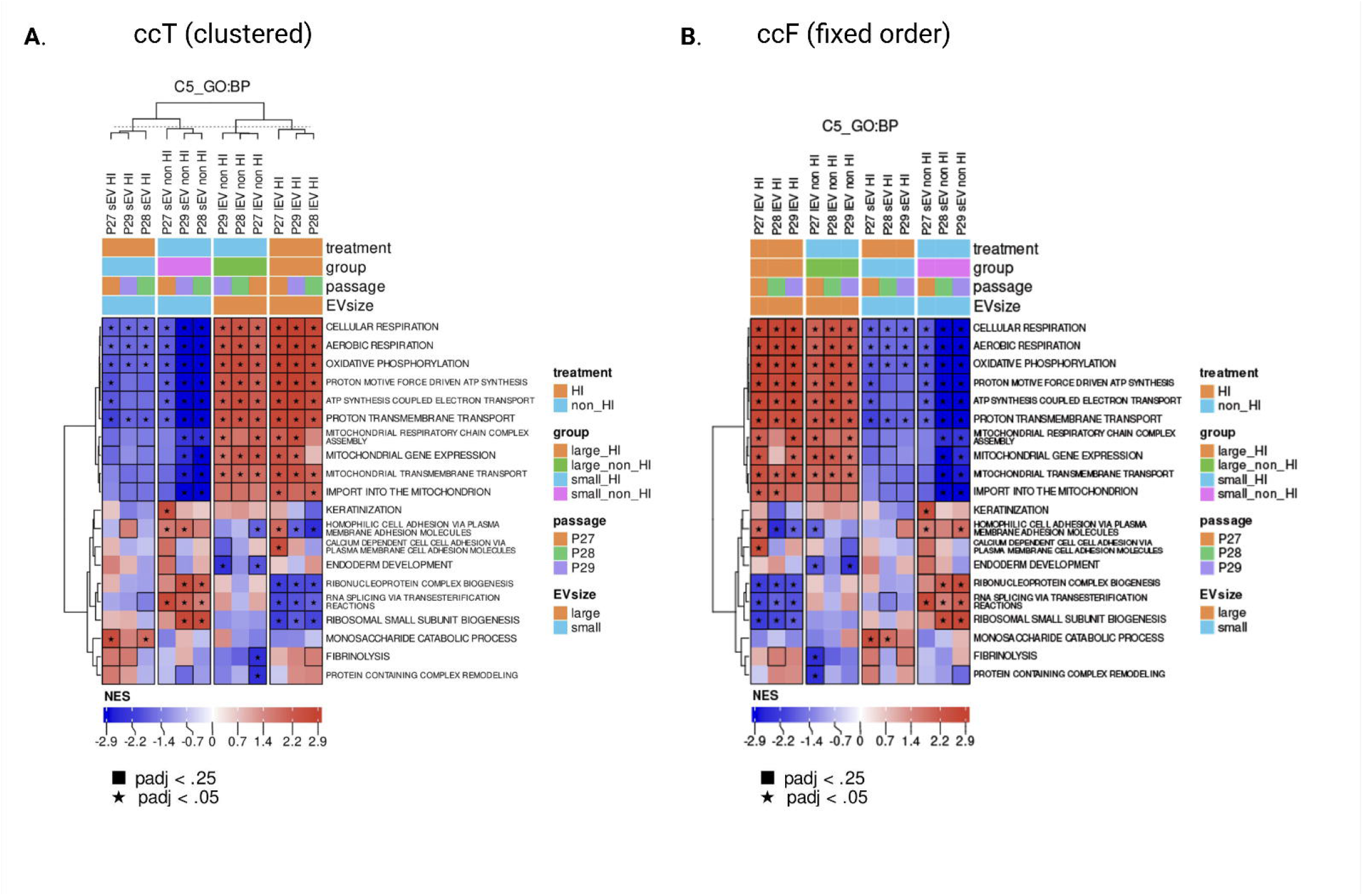
Mitochondrial and bioenergetic processes are enriched in large EVs. Per-sample gene set enrichment: normalized enrichment scores (**NES**) for the top 20 MSigDB C5 GO Biological Process gene sets in each of 12 EV samples (lEV and sEV; HI and non-HI serum; passages 27-29). (**A**) Samples clustered hierarchically (unsupervised). (**B**) Samples in fixed order, grouped by EV size and serum condition. Gene sets are clustered in both panels. **Red**, positive NES; **blue**, negative NES. Annotation bars indicate serum treatment, group, passage, and EV size. Star, adjusted p < 0.05; outlined cell, adjusted p < 0.25.

The most strongly and consistently enriched processes were mitochondrial. Oxidative phosphorylation showed the largest separation (mean NES + 2.64 in lEVs *vs*. -2.40 in sEVs; adjusted for p < 0.05 in all six EV samples), followed by cellular and aerobic respiration, ATP synthesis coupled electron transport, and proton transmembrane transport (**Fig. 1A, 1B; Supplementary Table S1**). Leading-edge proteins were canonical respiratory-chain and ATP synthase subunits, which included the following: Ndufs1, Ndufs3, Sdhb, Uqcrc1, Cyc1, Cox4i1, and Atp5f1d. Large EVs were also enriched in mitochondrial translation (mitoribosomal proteins such as Mrpl12 and Mrps6), mitochondrial protein import (Tomm40, Tomm22, Timm44, Samm50), fatty-acid beta-oxidation, and branched-chain amino-acid catabolism, indicating that lEV cargo spans both the respiratory chain and the pathways that supply it. As such, sEV-enriched processes, including ribosome biogenesis, protocadherin-mediated adhesion, basement-membrane organization, and glycolysis, were weaker and significant in only one to three of six sEV samples (**Supplementary Table S1**); we therefore regard them as secondary patterns.

Furthermore, running GSEA directly on the lEV *vs*. sEV comparison under each serum condition reproduced the per-sample findings as formal two-group tests: in both comparisons, mitochondrial and bioenergetic sets, including cellular and aerobic respiration and oxidative phosphorylation, were enriched in lEVs (**Figure 2**). The pattern was the same whether the top 10, 20, 40, or 80 gene sets were displayed (**Figure 2A-D**), indicating that it does not depend on gene-set selection depth.

**Figure 2.**
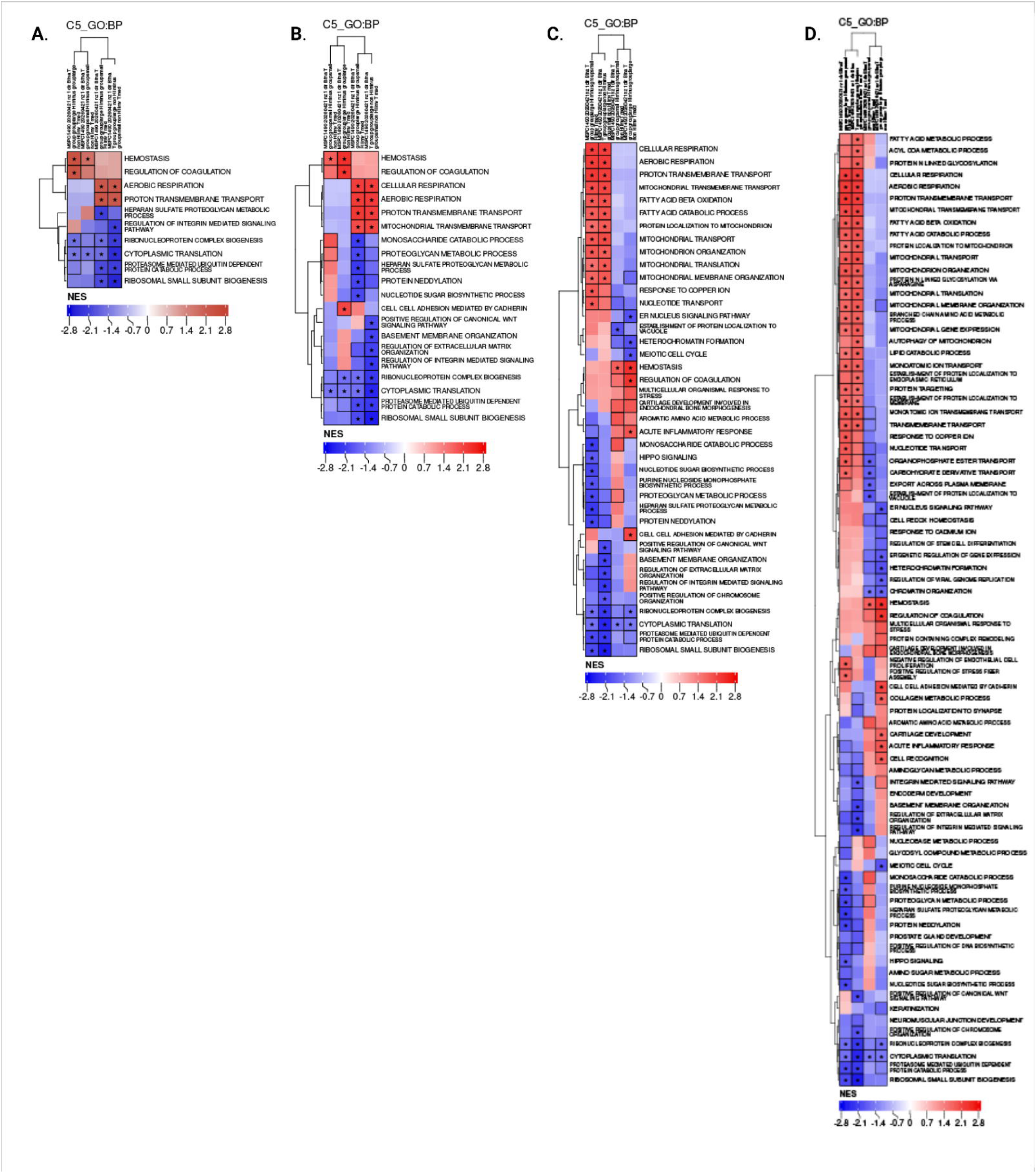
Per-comparison enrichment confirms mitochondrial enrichment in large EVs at every display depth. NES from pre-ranked GSEA of MSigDB C5 GO Biological Process gene sets for lEVs vs. sEVs under HI and under non-HI serum, with proteins ranked by signed log p-value and comparisons shown in fixed order, for the top (A) 10, (B) 20, (C) 40, and (D) 80 gene sets. Red, enriched in lEVs; blue, enriched in sEVs. Star, adjusted p < 0.05; outlined cell, adjusted p < 0.25. The two remaining columns compare HI with non-HI serum within lEVs and within sEVs (red, higher under HI serum); mitochondrial gene sets did not differ between serum conditions in either fraction.

## Discussion

The major finding of this study is that brain endothelial cell-derived lEVs are selectively enriched in mitochondrial and oxidative phosphorylation machinery relative to sEVs, and this enrichment is reproducible across passages and serum conditions. Enrichment of cellular respiration, oxidative phosphorylation, ATP synthesis, mitochondrial gene expression, and mitochondrial protein import in lEVs identifies vesicle size as the primary determinant of mitochondrial cargo (**Figure 1A, 1B**), and the same processes were enriched in lEVs in formal lEV *vs*. sEV comparisons under both serum conditions and at every display depth (**Figure 2**).

The co-enrichment of membrane-embedded respiratory complexes, outer-membrane import and assembly proteins, mitoribosomes, and fuel-oxidation enzymes suggests that lEVs carry membrane-bound mitochondrial material rather than a random sample of the cellular proteome (**Figure 1A, 1B**). These results are consistent with, and extend to endothelial-derived vesicles, the mitochondrial protein inventory that Cikic *et al*. described in cerebral microvessels, in which respiratory-chain subunits, VDAC channels, MICOS/SAMM50 assembly components, and ATP-synthase subunits constitute the core mitochondrial signature.^9^ These findings also agree with our prior direct evidence that BEC lEVs, but not sEVs, contain mitochondria visible by electron microscopy and transfer mitochondria to recipient cells.^5, 6^ We have also previously demonstrated that inhibiting respiratory complexes I and V in the donor BECs abolished the mitochondrial function transferred to recipient cells,^5, 8^ linking the respiratory machinery identified here to the functional activity of lEVs. They also converge with a growing literature showing that large EVs, but not small EVs, show selective effects of mitochondrial transfer in experimental models of cerebral ischemia.^5, 6, 8^

Moreover, the processes described above neatly fit canonical EV biogenesis pathways; lEVs bud from the plasma membrane and can incorporate mitochondria, whereas sEVs arise via endosomal pathways and are generally too small to package intact mitochondria.^10^ The co-enrichment of fatty-acid beta-oxidation and branched-chain amino-acid metabolism (**Table S1**) further suggests that lEVs carry a functionally coherent bioenergetic module rather than isolated respiratory components.

Small EV-enriched processes, including RNA splicing and processing, translation, and ribonucleoprotein complex biogenesis, homophilic cell adhesion, and monosaccharide catabolism, describe a cargo profile oriented toward RNA handling, surface interaction, and cytosolic rather than mitochondrial metabolism (**Figure 1A, 1B; Table S1**). This suggests that lEVs and sEVs carry distinct cargo programs rather than size variants of a common cargo, although we interpret the sEV signals cautiously given their smaller effect sizes and limited per-sample significance. For therapeutic development, a mitochondrial payload that is consistent across preparations is advantageous, since mitochondria-containing BEC EVs reduce injury in stroke models and their efficacy depends on the donor source.^4, 6^ Because the lEV mitochondrial signature was maintained under both serum conditions (**Figure 2**), mitochondrial cargo appears to be a stable property of the large-vesicle fraction. Replicates were preparations from consecutive passages (P27-P29).

## Conclusions

Our findings reveal that mitochondrial and bioenergetic machinery is selectively and reproducibly enriched in BEC lEVs relative to sEVs, strongly supporting the continued development of lEVs as novel carriers for mitochondria delivery. These findings also provide the first mechanistic insights for the bioenergetic effects of BEC-EV-mediated mitochondrial transfer.

## Acknowledgements

The authors gratefully acknowledge National Institutes of Health grant number 7R01NS136752-02 for funding support. We thank Mr. Balaji Govindaswamy for assistance with EV isolation and characterization. We acknowledge the services from Baylor College of Medicine’s Mass Spectrometry Proteomics Core led by Dr. Anna Malovannaya and Mr. Antrix Jain along with the support of Drs. Kwangwon Lee, Mei Leng, and Alexander Saltzman and Ms. Shirley Wang. The core is supported by CPRIT Core Facility Award (RP210227), P30 Cancer Center Support Grant (NCICA125123), Intellectual Developmental Disabilities Research Center Award (P50 HD103555), Gulf Coast Center for Precision Environmental Health (P30 ES030285), and S10 High End Instrument Award (OD026804).

## Author contributions

**Nikhila Achanta** designed methodology, completed the proteomic data analysis and wrote the manuscript. **Devika S Manickam** designed the study, analyzed the proteomics data and co-wrote and edited the manuscript.

