## Supplemental File for "Mitochondrial and bioenergetic processes are enriched in large- but are reciprocally depleted in small extracellular vesicles"

### Slide 1
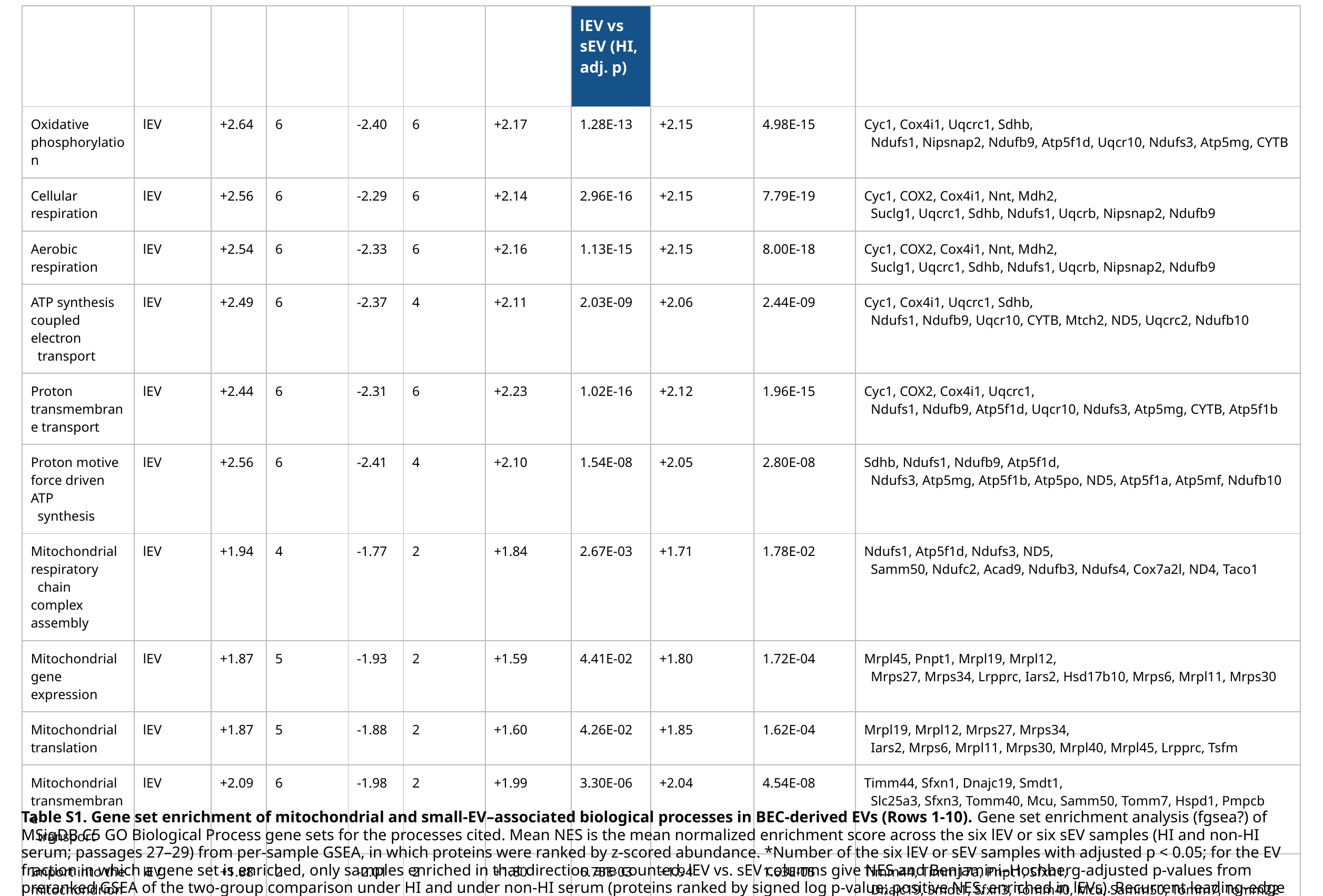

| Gene Set (GO:BP) | Enriched in | Mean NES (lEV) | lEV samples\* | Mean NES (sEV) | sEV samples\* | lEV vs sEV (HI NES) | lEV vs sEV (HI, adj. p) | lEV vs sEV (non-HI NES) | lEV vs sEV (non-HI, adj. p) | Recurrent leading-edge proteins (>/= 3 of 6 samples) |
| --- | --- | --- | --- | --- | --- | --- | --- | --- | --- | --- |
| Oxidative phosphorylation | lEV | +2.64 | 6 | -2.40 | 6 | +2.17 | 1.28E-13 | +2.15 | 4.98E-15 | Cyc1, Cox4i1, Uqcrc1, Sdhb, Ndufs1, Nipsnap2, Ndufb9, Atp5f1d, Uqcr10, Ndufs3, Atp5mg, CYTB |
| Cellular respiration | lEV | +2.56 | 6 | -2.29 | 6 | +2.14 | 2.96E-16 | +2.15 | 7.79E-19 | Cyc1, COX2, Cox4i1, Nnt, Mdh2, Suclg1, Uqcrc1, Sdhb, Ndufs1, Uqcrb, Nipsnap2, Ndufb9 |
| Aerobic respiration | lEV | +2.54 | 6 | -2.33 | 6 | +2.16 | 1.13E-15 | +2.15 | 8.00E-18 | Cyc1, COX2, Cox4i1, Nnt, Mdh2, Suclg1, Uqcrc1, Sdhb, Ndufs1, Uqcrb, Nipsnap2, Ndufb9 |
| ATP synthesis coupled electron transport | lEV | +2.49 | 6 | -2.37 | 4 | +2.11 | 2.03E-09 | +2.06 | 2.44E-09 | Cyc1, Cox4i1, Uqcrc1, Sdhb, Ndufs1, Ndufb9, Uqcr10, CYTB, Mtch2, ND5, Uqcrc2, Ndufb10 |
| Proton transmembrane transport | lEV | +2.44 | 6 | -2.31 | 6 | +2.23 | 1.02E-16 | +2.12 | 1.96E-15 | Cyc1, COX2, Cox4i1, Uqcrc1, Ndufs1, Ndufb9, Atp5f1d, Uqcr10, Ndufs3, Atp5mg, CYTB, Atp5f1b |
| Proton motive force driven ATP synthesis | lEV | +2.56 | 6 | -2.41 | 4 | +2.10 | 1.54E-08 | +2.05 | 2.80E-08 | Sdhb, Ndufs1, Ndufb9, Atp5f1d, Ndufs3, Atp5mg, Atp5f1b, Atp5po, ND5, Atp5f1a, Atp5mf, Ndufb10 |
| Mitochondrial respiratory chain complex assembly | lEV | +1.94 | 4 | -1.77 | 2 | +1.84 | 2.67E-03 | +1.71 | 1.78E-02 | Ndufs1, Atp5f1d, Ndufs3, ND5, Samm50, Ndufc2, Acad9, Ndufb3, Ndufs4, Cox7a2l, ND4, Taco1 |
| Mitochondrial gene expression | lEV | +1.87 | 5 | -1.93 | 2 | +1.59 | 4.41E-02 | +1.80 | 1.72E-04 | Mrpl45, Pnpt1, Mrpl19, Mrpl12, Mrps27, Mrps34, Lrpprc, Iars2, Hsd17b10, Mrps6, Mrpl11, Mrps30 |
| Mitochondrial translation | lEV | +1.87 | 5 | -1.88 | 2 | +1.60 | 4.26E-02 | +1.85 | 1.62E-04 | Mrpl19, Mrpl12, Mrps27, Mrps34, Iars2, Mrps6, Mrpl11, Mrps30, Mrpl40, Mrpl45, Lrpprc, Tsfm |
| Mitochondrial transmembrane transport | lEV | +2.09 | 6 | -1.98 | 2 | +1.99 | 3.30E-06 | +2.04 | 4.54E-08 | Timm44, Sfxn1, Dnajc19, Smdt1, Slc25a3, Sfxn3, Tomm40, Mcu, Samm50, Tomm7, Hspd1, Pmpcb |
| Import into the mitochondrion | lEV | +1.88 | 2 | -2.01 | 2 | +1.80 | 6.75E-03 | +1.94 | 1.63E-05 | Timm44, Timm17a, Pnpt1, Sfxn1, Dnajc19, Smdt1, Sfxn3, Tomm40, Mcu, Samm50, Tomm7, Tomm22 |
Table S1. Gene set enrichment of mitochondrial and small-EV–associated biological processes in BEC-derived EVs (Rows 1-10). Gene set enrichment analysis (fgsea?) of MSigDB C5 GO Biological Process gene sets for the processes cited. Mean NES is the mean normalized enrichment score across the six lEV or six sEV samples (HI and non-HI serum; passages 27–29) from per-sample GSEA, in which proteins were ranked by z-scored abundance. *Number of the six lEV or sEV samples with adjusted p < 0.05; for the EV fraction in which a gene set is enriched, only samples enriched in that direction are counted. lEV vs. sEV columns give NES and Benjamini–Hochberg-adjusted p-values from preranked GSEA of the two-group comparison under HI and under non-HI serum (proteins ranked by signed log p-value; positive NES, enriched in lEVs). Recurrent leading-edge proteins are those in the leading edge of at least three of the six samples of the enriched EV fraction. Cells labeled with 'n.t.' indicate gene sets not tested in that comparison.

### Slide 2
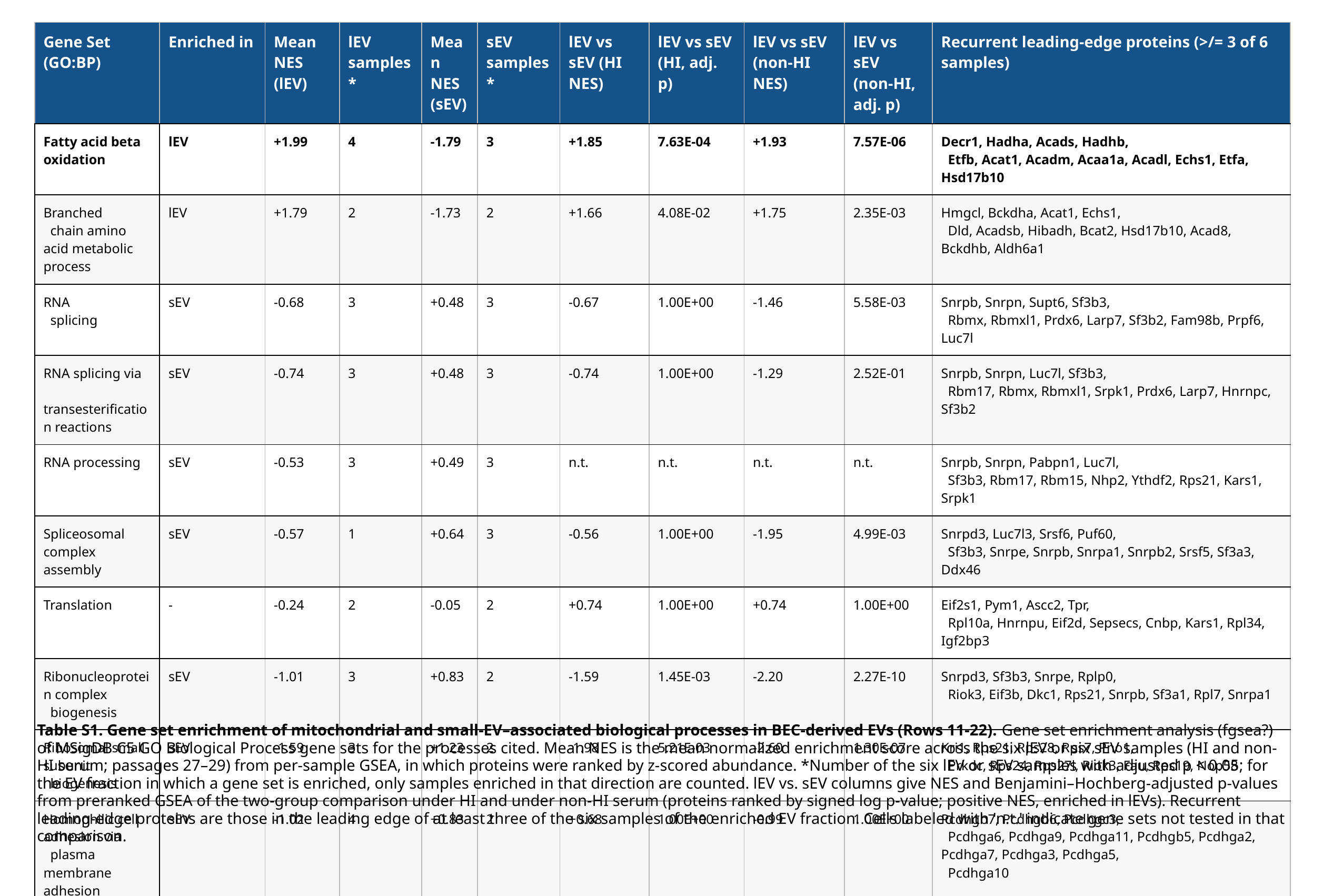

| Gene Set (GO:BP) | Enriched in | Mean NES (lEV) | lEV samples\* | Mean NES (sEV) | sEV samples\* | lEV vs sEV (HI NES) | lEV vs sEV (HI, adj. p) | lEV vs sEV (non-HI NES) | lEV vs sEV (non-HI, adj. p) | Recurrent leading-edge proteins (>/= 3 of 6 samples) |
| --- | --- | --- | --- | --- | --- | --- | --- | --- | --- | --- |
| Fatty acid beta oxidation | lEV | +1.99 | 4 | -1.79 | 3 | +1.85 | 7.63E-04 | +1.93 | 7.57E-06 | Decr1, Hadha, Acads, Hadhb, Etfb, Acat1, Acadm, Acaa1a, Acadl, Echs1, Etfa, Hsd17b10 |
| Branched chain amino acid metabolic process | lEV | +1.79 | 2 | -1.73 | 2 | +1.66 | 4.08E-02 | +1.75 | 2.35E-03 | Hmgcl, Bckdha, Acat1, Echs1, Dld, Acadsb, Hibadh, Bcat2, Hsd17b10, Acad8, Bckdhb, Aldh6a1 |
| RNA splicing | sEV | -0.68 | 3 | +0.48 | 3 | -0.67 | 1.00E+00 | -1.46 | 5.58E-03 | Snrpb, Snrpn, Supt6, Sf3b3, Rbmx, Rbmxl1, Prdx6, Larp7, Sf3b2, Fam98b, Prpf6, Luc7l |
| RNA splicing via transesterification reactions | sEV | -0.74 | 3 | +0.48 | 3 | -0.74 | 1.00E+00 | -1.29 | 2.52E-01 | Snrpb, Snrpn, Luc7l, Sf3b3, Rbm17, Rbmx, Rbmxl1, Srpk1, Prdx6, Larp7, Hnrnpc, Sf3b2 |
| RNA processing | sEV | -0.53 | 3 | +0.49 | 3 | n.t. | n.t. | n.t. | n.t. | Snrpb, Snrpn, Pabpn1, Luc7l, Sf3b3, Rbm17, Rbm15, Nhp2, Ythdf2, Rps21, Kars1, Srpk1 |
| Spliceosomal complex assembly | sEV | -0.57 | 1 | +0.64 | 3 | -0.56 | 1.00E+00 | -1.95 | 4.99E-03 | Snrpd3, Luc7l3, Srsf6, Puf60, Sf3b3, Snrpe, Snrpb, Snrpa1, Snrpb2, Srsf5, Sf3a3, Ddx46 |
| Translation | - | -0.24 | 2 | -0.05 | 2 | +0.74 | 1.00E+00 | +0.74 | 1.00E+00 | Eif2s1, Pym1, Ascc2, Tpr, Rpl10a, Hnrnpu, Eif2d, Sepsecs, Cnbp, Kars1, Rpl34, Igf2bp3 |
| Ribonucleoprotein complex biogenesis | sEV | -1.01 | 3 | +0.83 | 2 | -1.59 | 1.45E-03 | -2.20 | 2.27E-10 | Snrpd3, Sf3b3, Snrpe, Rplp0, Riok3, Eif3b, Dkc1, Rps21, Snrpb, Sf3a1, Rpl7, Snrpa1 |
| Ribosomal small subunit biogenesis | sEV | -1.59 | 3 | +1.23 | 2 | -1.98 | 5.21E-03 | -2.60 | 1.30E-07 | Kri1, Rps21, Rps28, Rps7, Pno1, Prkdc, Rps24, Rps27l, Riok3, Fau, Rps19, Nop58 |
| Homophilic cell adhesion via plasma membrane adhesion molecules | sEV | -1.02 | 4 | +0.83 | 2 | +0.68 | 1.00E+00 | -0.99 | 1.00E+00 | Pcdhgb7, Pcdhgb6, Pcdhgc3, Pcdhga6, Pcdhga9, Pcdhga11, Pcdhgb5, Pcdhga2, Pcdhga7, Pcdhga3, Pcdhga5, Pcdhga10 |
| Monosaccharide catabolic process | sEV | -0.73 | 0 | +0.79 | 2 | -2.13 | 4.08E-03 | -1.20 | 7.56E-01 | Pfkl, Eno1, Galk1, Hk1, Galt, Pgk1, Aldob, Pgm1, Eno3, Galm, Tkfc |
Table S1. Gene set enrichment of mitochondrial and small-EV–associated biological processes in BEC-derived EVs (Rows 11-22). Gene set enrichment analysis (fgsea?) of MSigDB C5 GO Biological Process gene sets for the processes cited. Mean NES is the mean normalized enrichment score across the six lEV or six sEV samples (HI and non-HI serum; passages 27–29) from per-sample GSEA, in which proteins were ranked by z-scored abundance. *Number of the six lEV or sEV samples with adjusted p < 0.05; for the EV fraction in which a gene set is enriched, only samples enriched in that direction are counted. lEV vs. sEV columns give NES and Benjamini–Hochberg-adjusted p-values from preranked GSEA of the two-group comparison under HI and under non-HI serum (proteins ranked by signed log p-value; positive NES, enriched in lEVs). Recurrent leading-edge proteins are those in the leading edge of at least three of the six samples of the enriched EV fraction. Cells labeled with 'n.t.' indicate gene sets not tested in that comparison.

### Slide 3
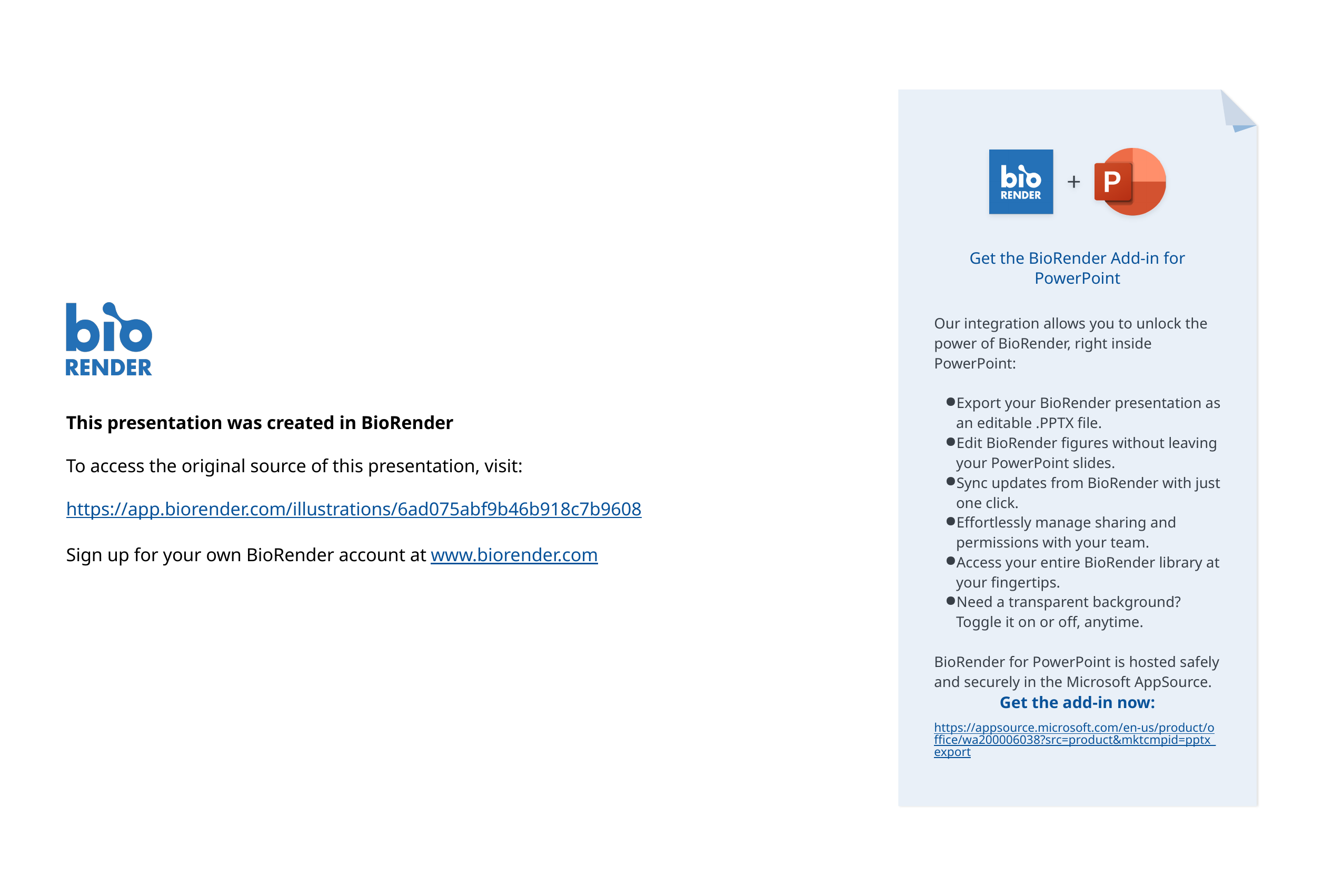

Get the BioRender Add-in for PowerPoint
Our integration allows you to unlock the power of BioRender, right inside PowerPoint:
Export your BioRender presentation as an editable .PPTX file.
Edit BioRender figures without leaving your PowerPoint slides.
Sync updates from BioRender with just one click.
Effortlessly manage sharing and permissions with your team.
Access your entire BioRender library at your fingertips.
Need a transparent background? Toggle it on or off, anytime.
BioRender for PowerPoint is hosted safely and securely in the Microsoft AppSource.
This presentation was created in BioRender
To access the original source of this presentation, visit:
https://app.biorender.com/illustrations/6ad075abf9b46b918c7b9608
Sign up for your own BioRender account at www.biorender.com
Get the add-in now:
https://appsource.microsoft.com/en-us/product/office/wa200006038?src=product&mktcmpid=pptx_export
